# Female immunity protects from cutaneous squamous cell carcinoma

**DOI:** 10.1101/2021.01.28.428489

**Authors:** Timothy Budden, Caroline Gaudy-Marqueste, Sarah Craig, Yuan Hu, Charles Earnshaw, Shilpa Gurung, Amelle Ra, Victoria Akhras, Patrick Shenjere, Ruth Green, Lynne Jamieson, John Lear, Luisa Motta, Carlos Caulín, Deemesh Oudit, Simon J Furney, Amaya Virós

## Abstract

**Purpose:** Cancer susceptibility and mortality are higher in males, and the mutational and transcriptomic landscape of cancer differs by sex. The current assumption is that men are at higher risk of epithelial cancers as they expose more to carcinogens and accumulate more damage than women. We present data showing women are more protected from aggressive cutaneous squamous cell carcinoma (cSCC) due to strong immune activation.

**Methods:** We explored clinical and molecular sexual disparity in immunocompetent and immunosuppressed patients (N= 738, N=160) with carcinoma cSCC, in FVB/N mice exposed to equal doses of DMBA, and in human keratinocytes by whole exome sequencing, bulk and single cell RNA sequencing.

**Results:** We show cSCC is more aggressive in men, and immunocompetent women develop mild cSCC, later in life. To test if sex drives disparity, we exposed male and female mice to equal doses of carcinogen, and found males present more aggressive, metastatic cSCC than females. Critically, females activate cancer immune-related expression pathways and CD4 and CD8 T cell infiltration independently of mutations. In contrast, males increase the rate of mitoses and proliferation in response to carcinogen. Human female skin and keratinocytes also activate immune-cancer fighting pathways and immune cells at ultraviolet radiation-damaged sites. Critically, a compromised immune system leads to high-risk, aggressive cSCC specifically in women.

**Conclusions:** This work shows the immune response is sex biased in cSCC, and highlights female immunity offers greater protection than male immunity.

## Introduction

Many diseases show sex disparity in their epidemiology, clinical course and outcome(1), and cancer incidence is higher in men even after adjusting for risk factors. Cancer mortality also affects men disproportionately and women respond better to some cancer therapies(2–4). Squamous cell carcinomas (SCCs) arise in epithelia from the head and neck, lung, bladder, esophagus and skin, have a strong male bias and are primarily caused by carcinogens such as tobacco, alcohol and ultraviolet radiation (UVR). The higher SCC male incidence and mortality is thought to reflect the increased lifetime exposure of men to carcinogens and delayed clinical care; and molecular studies show a higher burden of carcinogen-driven mutations in male tumors(5–9). SCCs arising on the skin, cutaneous SCC (cSCC)(10–12), are more frequent in men, the 2^nd^ most frequent malignancy in humans and the most common malignancy in patients with a compromised immune system(10,13,14). In this study, we examined if the higher incidence and mortality of cSCC in men is due to increased vulnerability to epithelial neoplasia in the male sex, rather than due to increased exposure to a carcinogen. To test this hypothesis, we used the most frequent cSCC mouse model driven by the topical cutaneous application of DMBA, a polycyclic aromatic hydrocarbon that induces epidermal *Hras* mutations, followed by topical tetradecanoyl-phorbol acetate (TPA), which induces inflammation and epidermal proliferation, leading to epidermal papillomas and cSCC(15). The advantages of this model are that the genomic landscape of DMBA/TPA-induced cSCC displays significant overlap with human cutaneous SCC, and that it allows the *in vivo* study of cSCC controlling for age, strain, susceptibility and dose of carcinogen(16). We explored the clinical and molecular sex bias of cSCC carcinogenesis in animals exposed to the same dose of carcinogen, and confirmed the molecular findings in human skin and keratinocytes. Additionally, we examined the relationship between age at diagnosis, histological grade of cSCC and immune status in human cohorts to explore the role of immunity by sex.

## Methods

### Animal experiments

Experiments were performed in 4-week-old FVB/N male and female mice, and DMBA (25mg/ml) and TPA (0.02mg/ml) in acetone applied once a week, two days apart for 6 weeks, followed by TPA weekly for 10 weeks or until tumor development.

### Molecular analyses

DNA from the largest sized mouse tumor, adjacent treated skin and a kidney was sequenced, patterns of single nucleotide variation, insertion and deletions in tumors and skin were analysed and oncoplots generated of the top 20 frequently mutated genes by sex and histology to identify patterns of mutations. Paired-end RNA sequencing was performed from fresh whole DMBA/TPA treated skin from the back and normal skin from the abdomen. Human single cell RNA-seq data from male and female keratinocytes was analysed from a study of squamous cell carcinoma(17), downloaded from GEO database (GSE144236).

### Histology

Mouse tumors were classified as papillomas, keratoacanthoma, well, moderately and poorly differentiated cSCC, and visceral organs stained with H&E and pan-keratin to confirm metastasis. Immunohistochemistry of tumors and skin was performed with anti-phospho-Histone H3 (Ser10) (06-570) CD4 Antibody (14-9766-82) ThermoFisher / eBioscience and CD8a Antibody (14-0808-82) ThermoFisher / eBioscience.

### Human samples analysis

Age, sex, immune status and histological grade of human primary cSCC from 3 UK NHS hospitals from immune-competent and immune-suppressed patients diagnosed by NHS pathologists under routine diagnostic practice were collected. cSCC were classified as keratoacanthoma (KA)/ well differentiated invasive SCC (1), moderately differentiated SCC (2) and poorly differentiated SCC (3). The immune status of patients was retrieved from clinical histories, and immunosuppressed patients were organ transplant recipients on immunosuppressive medication, had white blood cell dyscrasia or systemic cancer treatment with immunotherapy, radiotherapy or chemotherapy in the past 10 years, chronic inflammatory disorders and autoimmune disorders on systemic immunosuppressive medication.

## Results

### Immune competent men develop more aggressive cSCC than women

To explore if men are more susceptible to aggressive cSCC than women, we studied the relationship between sex, age and histological grade in consecutive cSCC samples excised from immunocompetent patients (n = 738; men: 459, women: 279, Supplementary Table 1). We confirmed cSCC more commonly affects immunocompetent men (62.2% men), and found the median age of diagnosis to be similar in men and women (median age men = 79, women = 81, p=0.10, Fig. 1A). Strikingly, men more frequently presented cSCC that was more aggressive, less differentiated, of higher histological grade compared to cSCC of women (Fig. 1B, p=0.0002). Furthermore, when we restricted our study by sex to the more aggressive variants of cSCC, which have the highest risk of metastasis(14), we found that immunocompetent females diagnosed with higher grade cSCC were older than men (median age men =80, median women =83, p=0.04, Fig. 1C). These data suggest that men are at higher risk of aggressive cSCC.

**Figure 1.**
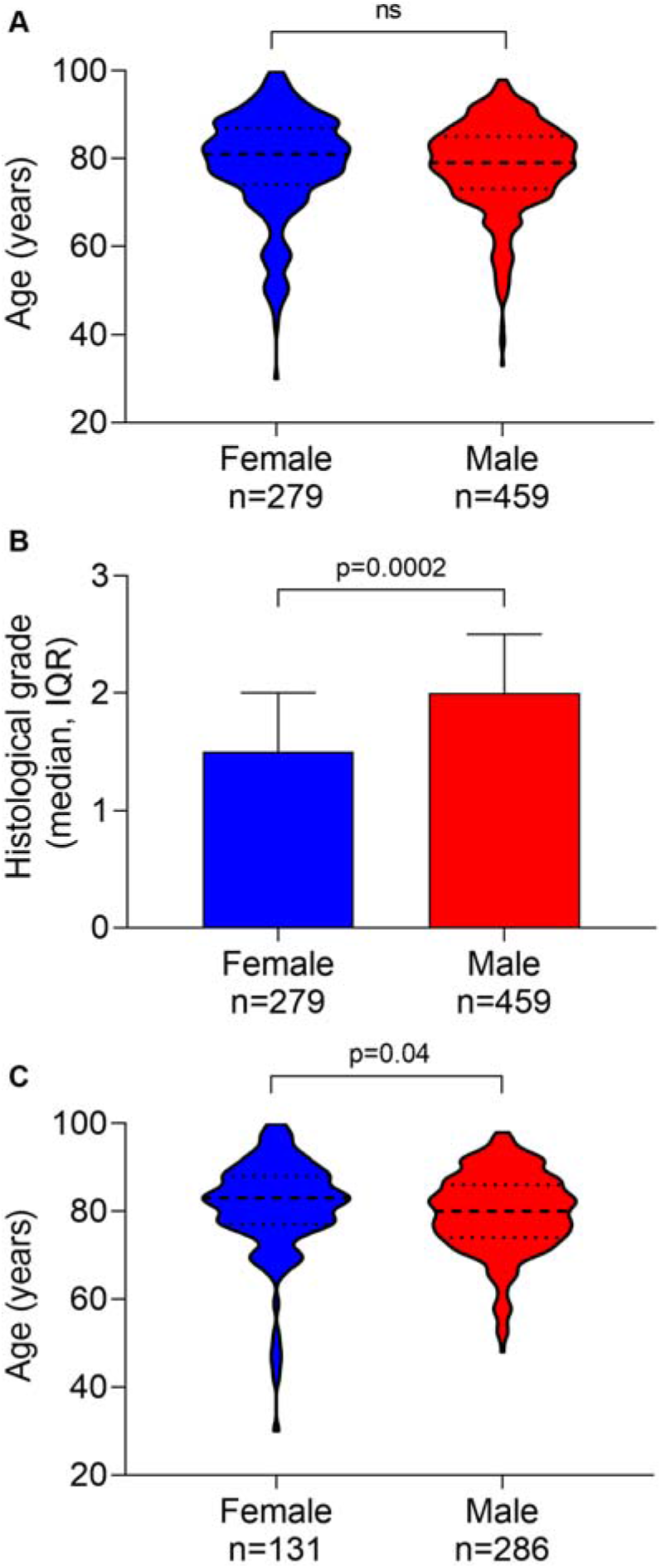
Men have more aggressive cSCC. (A) Age of immunocompetent patients diagnosed with primary cSCC by sex. (B) Histological grade of primary cSCC by sex in immunocompetent patients (median, interquartile range (IQR). (C) Age of immunocompetent women and men diagnosed with moderately and poorly differentiated primary cSCC.

### Male mice are more susceptible to chemically-induced aggressive cSCC

To determine if the increased susceptibility to aggressive cSCC in men is due to a greater exposure to carcinogens or due to biological sex differences, we exposed immunocompetent male and female mice, matched for age and strain, to equal doses of the carcinogen DMBA/TPA, which promotes cSCC in mice. We recorded earliest lesion incidence and burden, and found males developed more papillomas and cSCC earlier than female animals, although this difference was not statistically significant (p=0.10, Fig. 2A). Animals presented a range of squamoproliferative lesions, including epidermal hyperplasia, papillomas, well-differentiated, invasive SCC and more aggressive, spindle SCC. However, compared to females, males presented more advanced, aggressive subtypes of disease (Fig. 2B). Specifically, male cSCC presented more mesenchymal, spindle features, compared to female cSCC. Importantly, only males presented metastatic lung cSCC deposits (Fig. 2B, 2C). Taken together, these data show male mice developed more aggressive, metastatic cSCC than females exposed to the same dose of carcinogen.

**Figure 2.**
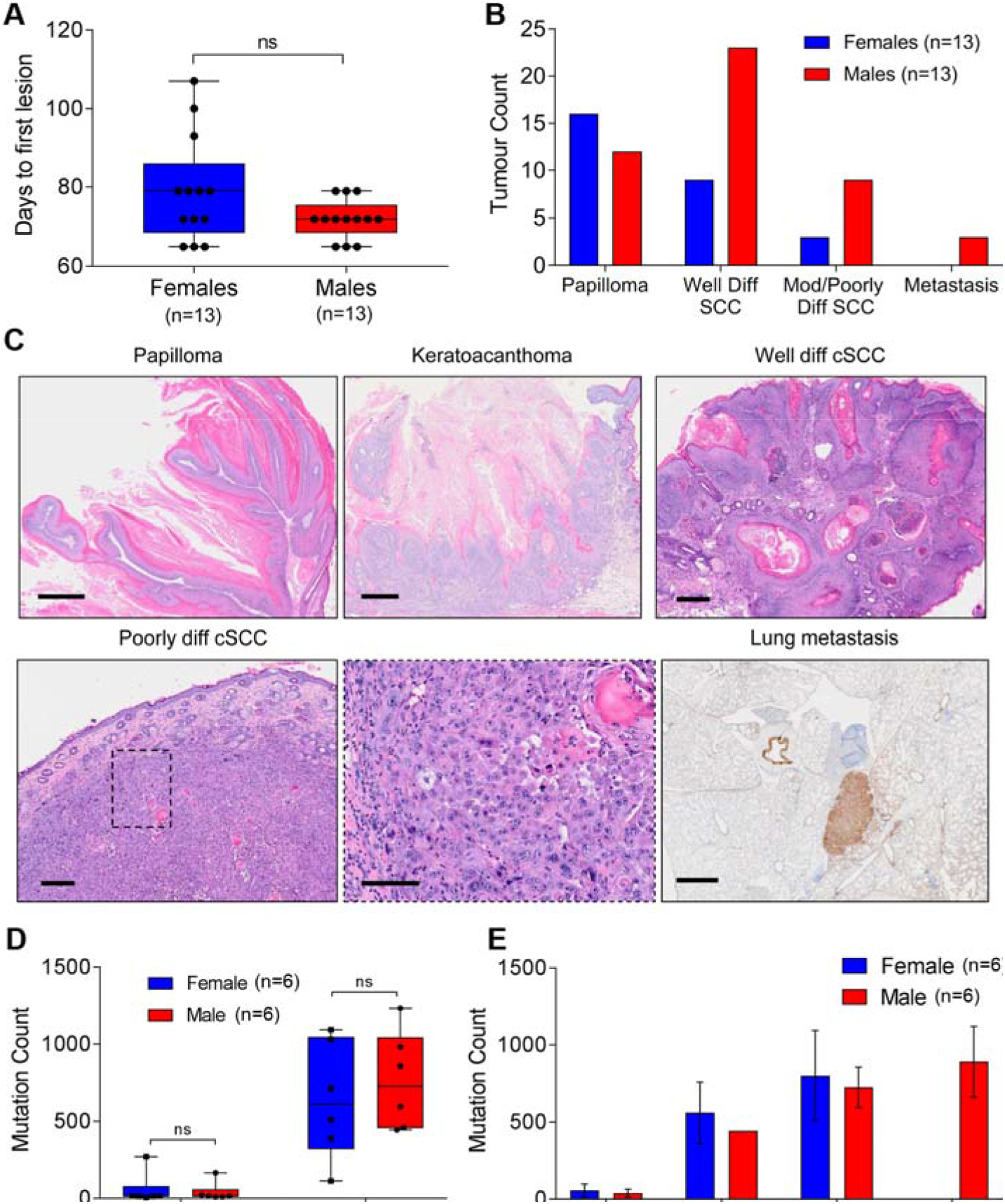
Male animals are more susceptible to chemically-induced aggressive cSCC. (A) Days to first lesion in male and female skin exposed to DMBA/TPA. (B) Histological grade of cSCC tumors by sex. (C) Representative H&E of papilloma (scale bar: 400μm), keratoacanthoma (scale bar: 500μm), well differentiated cSCC (scale bar: 500μm), poorly differentiated cSCC (scale bar: 500μm) and insert (box, scale bar: 100μm) and lung metastasis stain for pan-cytokeratin (brown, scale bar: 1mm). (D) Tumor mutation burden in DMBA/TPA exposed skin (Treated Skin) and DMBA/TPA-induced cSCC (Tumor) by sex (E) Tumor mutation burden in DMBA/TPA lesions by histological grade and sex. Error bars represent standard error of the mean (bar).

### DNA damage accumulates equally in male and female animal skin and cSCC

Increased mutation burden underpins more aggressive forms of epithelial cancer(18), and human male epithelial tumors have a higher mutation burden(5–9). Therefore, we examined if the more aggressive male cSCCs in our animals are due to greater mutation accumulation, or less DNA repair, leading to higher mutation burden. For this, we compared the tumor mutation burden (TMB) in DMBA/TPA-induced mouse cSCCs (DT-cSCC), and found deep targeted exome sequencing of DT-cSCC revealed the number of total mutations increased with increasing histological grade, but did not differ by sex (Fig. 2D, 2E). Furthermore, we found no differences in the pattern and types of mutations by sex (Supplementary Table 2, Supplementary Fig. 1A).

Mouse tumors display a range of morphological features within the same histological grade, and cancer-driving mutations can associate specific histological features(19). Therefore, to ensure specimens were exactly comparable between males and females, we focused on mutations accrued in clinically and histologically normal DMBA/TPA-treated skin (DT-skin). We found males and females accumulate the same amount of genetic damage in DT-skin (Fig. 2D), and similar to DT-cSCCs, there was no sex disparity in the number, pattern and types of mutations (Supplementary Fig. 1B, Supplementary Table 2). These data indicate carcinogens damage male and female DNA similarly, and the more aggressive phenotype of male cSCC is independent of mutation burden.

### Unique transcriptomic and immune changes by sex in mice

Male animals develop more aggressive cSCC independently of the mutation spectrum. Therefore, we explored if the transcriptomic response to carcinogen exposure differed by sex. To reduce the gene expression variability due to histological differences between tumors, we compared the histologically normal, carcinogen exposed DT-skin of males and females. We first investigated the autosomal gene expression changes in DT-skin and normal skin, and then studied whether DT-skin gene expression varied by sex. We found the most significant changes in DT-skin involved critical cancer immune regulatory pathways including the interferon gamma and interferon alpha response pathways; as well as the inflammation-related and allograft rejection genes (Fig. 3A, Supplementary Table 3). In contrast, normal, untreated skin expressed genes involved in RAS signaling (Fig. 3A). We then investigated sex-specific changes in DT-skin, and found female DT-skin presented more transcriptomic changes than male skin overall, including critical genes involved in cancer immunity. We noted specifically the cytokine interferon gamma (*Ifng*), which is known to drive potent antitumor activity(20–22), to be increased in female carcinogen-treated skin. Intriguingly, we additionally observed female upregulation of the G1 to S-phase tumor suppressor, senescence-inducing cell cycle regulator *Cdkn2a* (Fig. 3B, 3C, Supplementary Table 4). Cdkn2a can exert ample anti-tumor effects(23,24), and when expressed in keratinocytes restricts proliferation, increases senescence and differentiation, and reduces tumor growth(25), consistent with its tumor-suppressive roles. Furthermore, recent work shows natural and immune checkpoint cancer immune control is achieved via Cdkn2a-dependent signaling, and interferons directly activate *Cdkn2a*(26).

**Figure 3.**
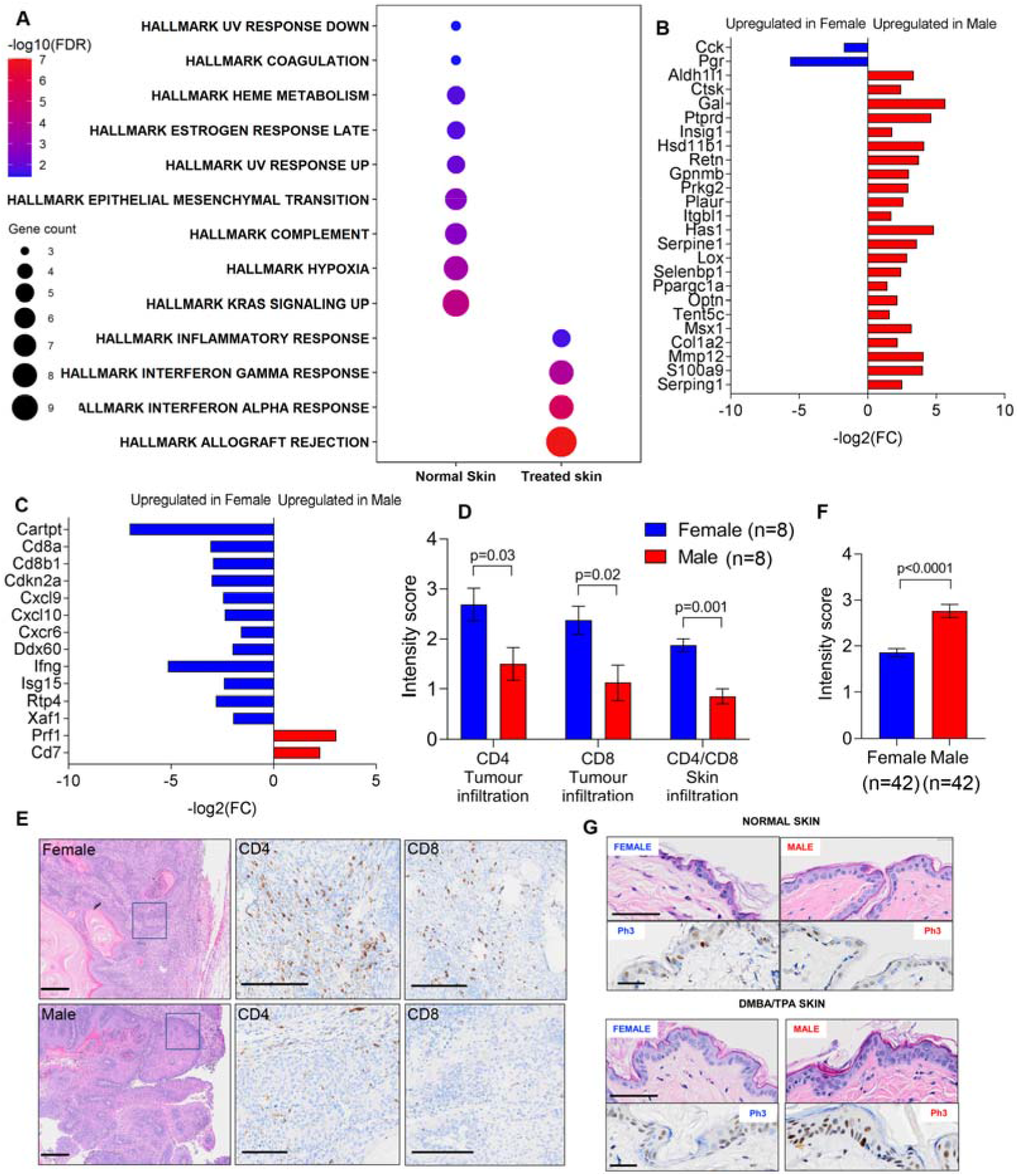
Female mouse skin upregulates immune cancer pathways and cells in response to chemical carcinogenesis. (A) Gene pathways differentially expressed by sex in normal skin and in DMBA/TPA-exposed skin (treated skin). (B) Differentially expressed genes enriched in pathways in normal skin and (C) treated skin. Blue: upregulated in female skin, red: upregulated in male skin. (D) Quantification of CD4 and CD8 in tumors and DMBA/TPA tumors and treated skin by sex. (E) Representative images of H&E and CD4 and CD8 T cell IHC in male and female cSCC (scale bar: 200μm). (F) Intensity score and (G) photomicrographs of mitotic cells (phospho-H3) by sex in DT-skin.

We next investigated if, in addition to immune-gene expression changes, female animals present a unique immune cell landscape. For this, we studied the proportion and subtypes of adaptive and innate immune cells by gene expression, which revealed a trend in female skin for increased antigen-presenting CD1 cells and CD4 / CD8 tumor lymphocytes. In contrast, male animal skin tended to be enriched for macrophages (Supplementary Fig 1C). We explored gene expression by immunohistochemistry staining of CD4 and CD8 T cells in DT-cSCC and DT-skin by sex, and confirmed female tumors and female DT-skin were more infiltrated with immune cells than male DT-cSCC and DT-skin (Fig. 3D, 3E).

Tumors in animals arise weeks after being exposed to the DMBA carcinogen, but the total number of mutations per cSCC rises progressively in tumors of increasing histological grade (Fig. 2E). We hypothesized the selection and growth of the more mutated clones leading to more aggressive cSCC occurs due to increased cell division and a proliferation advantage of these clones. As males present more aggressive disease, and female epidermis expresses higher levels of the cell cycle regulator *Cdkn2a*, we investigated if a higher epidermal proliferation rate in the male epidermis could underpin speedier clonal selection and disease progression of male disease. For this, we compared the rate of the mitosis-specific marker phospho-histone 3 (phospho-H3) in male and female epidermis and observed that DT-skin, compared to animal-matched non-DT skin was thickened, presenting increased number of layers (Supplementary Fig. 1D, p<0.0001). Next, we compared male to female DT-epidermis and found that phosphoH3 in male epidermis was increased in males (Fig. 3F, 3G, p<0.0001). We compared the expression of phospho-H3 in a subset of cSCCs, and again observed a trend for higher expression in males (Supplementary Fig. 1E, S1F). Thus, male and female mouse epidermis modulate the rate of epidermal proliferation differently following exposure to a chemical carcinogen.

Taken together, these findings indicate the transcriptome and immune cell response of male and female mice differs at the earliest stage of carcinogenesis, independently of DNA damage. Males present higher rates of epidermal proliferation following chemical carcinogen exposure. In contrast, females strongly upregulate immune responses and cancer-linked lymphocytes, implicating immunity in delayed female tumorigenesis.

### Human female skin upregulates immune-related cancer defense pathways

To validate the role of immune-related expression changes and immune cells modulating carcinogenesis in females, we compared the sex-specific changes in single cell RNA of normal male and female human keratinocytes obtained from cSCC-adjacent skin(17). Human cSCC is driven by ultraviolet light and arise in fields of severely sun-damaged skin, so we reasoned cSCC-adjacent keratinocytes, from sun exposed anatomic sites of adults, will accumulate significant carcinogen exposure. Indeed, we found that similar to animal DT-skin, the most prominently upregulated pathways in human keratinocytes overlapped with mouse skin. Strikingly, the most prominent pathway expression changes in all female keratinocyte subtypes were up-regulation of interferon gamma, interferon alpha and allograft rejection pathway genes (Fig. 4A, Supplementary Table 5-7). In further agreement with the mouse experiment, we found female human sun-exposed epidermis and cSCCs had a similar trend to higher counts of CD4, CD8 T cells and CD1C dendritic cells, whilst male skin and cSCC showed a trend for more macrophages (Supplementary Fig. 1G, 1H). Thus, these data indicate that the biological response to chemical and ultraviolet light carcinogen exposure in keratinocytes varies by sex in mice and humans.

**Figure 4.**
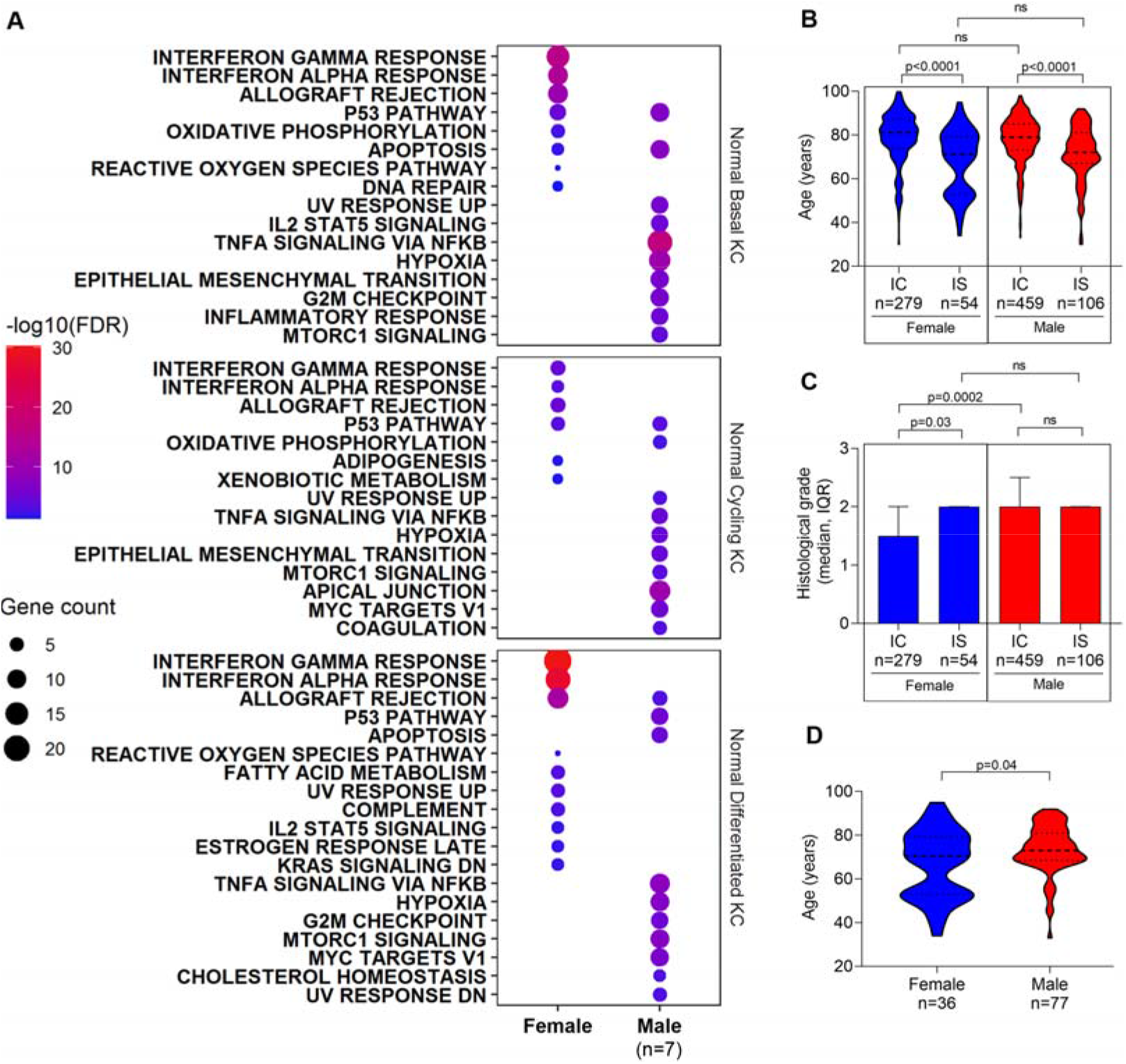
cSCC histological grade of human cSCC is tied to female immunity. (A) Pathways enriched for genes differentially expressed by sex in human adult keratinocytes from UVR-exposed, tumor-adjacent skin by sex. (B) Age comparison of immunocompetent (IC) and immunosuppressed (IS) patients diagnosed with primary cSCC by sex. (C) Histological grade comparison of primary cSCC by sex in immunocompetent (IC) and immunosuppressed (IS) patients (median, interquartile range (IQR). (D) Age comparison of immunocompetent (IC) and immunosuppressed (IS) women and men diagnosed with moderately and poorly differentiated primary cSCC.

### Female immunity inhibits aggressive cSCC

Because we found the immune response upregulated in female animals, and immune competent women have less aggressive forms of cSCC (Fig. 1B), we reasoned women with compromised immune systems would present more aggressive cSCC, comparable to immunocompetent men. To test this hypothesis, we investigated sex, age and histological grade in cSCC specimens excised from a cohort of immunosuppressed patients (IS, n= 160, men =106 and women n=54, Supplementary Table 1). Strikingly, we discovered cSCC appeared at a younger age in both immunosuppressed men and women, supporting the critical role of immunity in delaying cancer in both sexes (median age men =72, women =71, p=0.25, Fig. 4B). We then compared the histological grade in immunosuppressed and immunocompetent women, which revealed immunosuppressed women present more histologically aggressive disease (p=0.029, Fig. 4C), of equivalent grade to men. By contrast, men present aggressive disease regardless of their underlying immune status (Fig. 4C).

Next, we restricted our study by sex to the more aggressive, risky variants of cSCC in the immunosuppressed population, and found, as expected, that aggressive disease occurred earlier in life in both sexes. However, the lower age at diagnosis particularly occurred in women, suggesting the loss of immunity is more damaging to females (median age men =73, median women =70, p=0.04, Fig. 4D). Taken together, these data show the immune system is critical to contain carcinogenesis, and validate the immune response underpins less aggressive disease in females.

## Discussion

The molecular landscape of cancer differs by sex, and men are more frequently diagnosed with cancer and die more of cancer than women. Men traditionally are exposed to more carcinogens due to professional hazards and behavior, but in risk-adjusted epidemiological studies, there is still an unexplained greater proportion of men with cancer compared to women(2–4). Furthermore, there is significant variation in the DNA alterations and RNA landscape by sex in most cancers, suggesting there are sex-specific susceptibilities and biological processes driving female and male cancer(5–9). However, comparing human tumors from different sexes is a limited approach as human tumors are not matched for age, histological grade, underlying susceptibility to cancer, and dose of carcinogen.

In this study we show immunocompetent women have less aggressive cSCC than men. We compared the development and the molecular hallmarks of cSCC in male and female animals using a well-established mouse model of carcinogenesis, adjusted for carcinogen exposure(16). We show that the rate of DNA damage accumulation is equal in male and female mice exposed to equal doses of DMBA. This indicates both sexes repair damage similarly following exposure to the chemical carcinogen. Intriguingly, we observe the overall mutation burden (TMB) increases as tumors become more aggressive, or with increasing histological grade in both sexes; despite animals not receiving additional exposures to carcinogen. This suggests that in DMBA-driven cSCC there is a selection for clones with more mutations as cell division progresses and tumors advance. Previous work comparing mutation burden in adjacent areas of early dysplasia, intermediate dysplasia and primary cutaneous melanoma reveal increasing mutation burden with oncogenic stage(27); raising the hypothesis that areas with early damage must receive additional carcinogenic exposure to drive cancer progression. However, our study indicates carcinogenic damage conferring full oncogenic potential may occur at the early stage and will depend on successful expansion of clones with higher TMB, as we discontinued carcinogen exposure before cSCC onset.

Intriguingly, we find it is the transcriptional response to carcinogen exposure that differs between the sexes. At the earliest stage of disease, female animals increase cancer immune-related responses and recruit an immune cell landscape involved in cancer defense. Male animals, by contrast, have more macrophages, which are linked to poor prognosis in cancer, and comparatively show few immune-related gene expression changes. One critical gene expression difference we found between male and female animals is the up-regulation of *Cdkn2a* in females, a fundamental cell cycle regulator that exerts profound influence in both cell proliferation(23,24) and cancer-immune responses(25,26). Based on this finding, we explored the possible differences in the mitotic rate of carcinogen-exposed epithelium between males and females, and found male epidermis was more proliferative than female skin.

We validated the relevance of the *in vivo* findings in human female and male epidermal keratinocytes, which originated from a sun-exposed site(17), and in human cSCCs. Keratinocytes from women mirror the response observed in female mouse epidermis, up-regulating critical cancer-immune pathways, genes and immune cells following UVR. Moreover, like in the mouse model, although cSCC arises at a similar age in immunocompetent men and women, men have significantly more aggressive disease, and the aggressive tumors arise at a younger age in men. Restricting our analysis to immunosuppressed patients, we show cSCC arises in younger men and women. Importantly, cSCC in immunosuppressed women are significantly more aggressive, similar to male disease, whilst male disease is aggressive regardless of immune status. These data show the immune response plays a critical role constraining cSCC progression at the earliest stage of disease in both sexes, as the age of incidence drops in both men and women. However, the data indicate the mechanisms restricting tumor progression are more robust in females. Thus, this study, adjusted for risk factors, implicates distinct biology driving male and female cancer; and shows differences in immune responses between the sexes. These data are strongly aligned with clinical observations revealing higher incidence of excess immunity-linked disease in females and unique immune responses to infectious disease by sex(28). Personalized medicine approaches stratify cancer patients by genotype; however, to date, the potential for cancer stratification and therapy by sex has not been explored. Further work will be necessary to identify the potential regulators of sex-linked cSCC dimorphism.

## Acknowledgements

We are grateful to the Khavari Lab for advice and shared data.

## Author contributions

Conceptualization: A.V.; Writing of the original manuscript and generation of figures T.B. and A.V.; Collection of data: T.B., A.V., S.C., C.E., A.R., V.A., P.B., D.O., R.G., L.J., L.M., J.L.., H.Y., C.C.; Analysis of data: T.B., A.V., L.M., S.J.F., H.Y., C.C.; Tools and methods: T.B., A.V., S.J.F., C.C.; Software: T.B., S.J.F.; Funding acquisition, project administration, resources, supervision: A.V.

## Supplementary Material

**Supplementary Figure 1.**
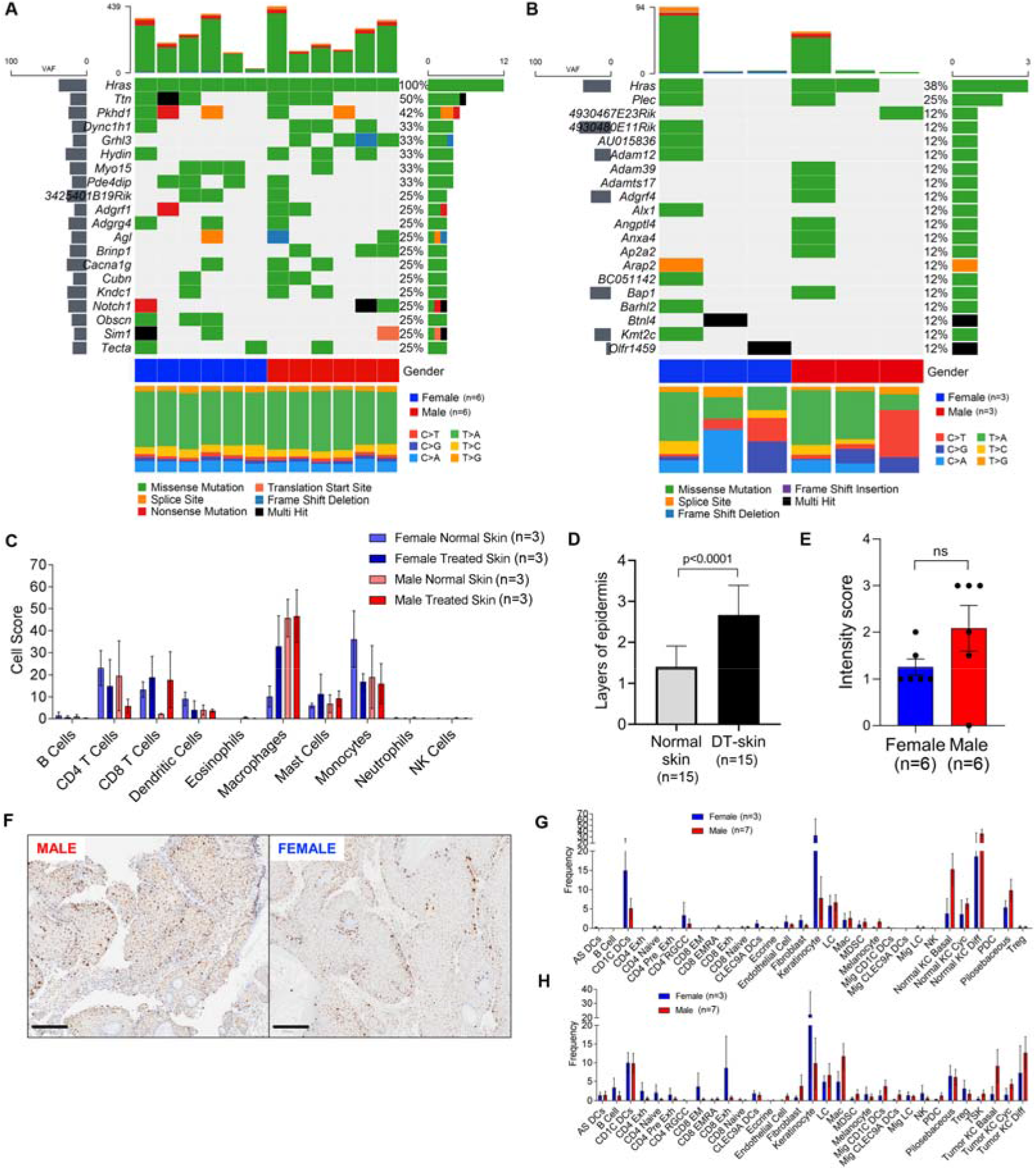
(A) Oncoplots of 20 most frequently mutated genes by sex in DT-cSCC and (B) DMBA/TPA treated skin (DT-skin). (C) Immune cell landscape in normal and DMBA/TPA treated skin by sex estimated from RNA-seq data(1), p values not significant, (n males = 3; females = 3) (D) Layers of epidermal cells in normal skin and DT-skin. (E) Intensity score of mitotic cells (phospho-H3) by sex in DT-cSCC. (F) Representative images of phospho-H3 staining (brown) in male and female DT-cSCC tumors (scale bar: 200μm). (G) Immune cell landscape in human adult UVR-skin and (H) human UVR-cSCC by sex p values not significant, (n males = 7; females = 3).

## Supplementary Tables

**Supplementary Table 1.** Clinical characteristics of 3 cSCC clinical cohorts

**Supplementary Table 2.** Mutations in tumors and DMBA/TPA treated skin

**Supplementary Table 3.** Differentially expressed genes by sex in normal skin

**Supplementary Table 4.** Differentially expressed genes by sex in DMBA/TPA treated skin

**Supplementary Table 5.** Genes differentially expressed by sex in normal tumor adjacent basal keratinocytes

**Supplementary Table 6.** Genes differentially expressed by sex in normal tumor adjacent cycling keratinocytes

**Supplementary Table 7.** Genes differentially expressed by sex in normal tumor adjacent differentiated keratinocytes

## Supplementary Methods

### Animal experiments

All procedures involving animals were performed under the Home Office approved project license P8ADED6C8, in accordance with ARRIVE guidelines and UK Home Office regulations under the Animals (Scientific Procedures) Act 1986. The study received ethical approval by the Cancer Research UK Manchester Institute’s Animal Welfare and Ethics Review Body (AWERB). All mice were maintained in pathogen-free, ventilated cages in the Biological Resources Unit at our Institute, and allowed free access to irradiated food and autoclaved water in a 12=h light/dark cycle, with room temperature at 21◻±◻2◻°C. All cages contained wood shavings, bedding and a cardboard tube for environmental enrichment. Experiments were performed in two cohorts of 4-week-old FVB/N mice (cohort 1: 6 females, 6 males, cohort 2: 7 females, 7 males). DMBA (25mg/ml) and TPA (0.02mg/ml) diluted in acetone was applied separately to the shaved backs of mice once a week, two days apart for 6 weeks, followed by TPA only weekly for a further 10 weeks or until first tumors started to develop. Animal tumor development was closely monitored and once tumor development began tumors were counted weekly and classified by size (papule: palpable 1-4mm, papilloma: exophytic, pedunculated lesions of small diameter 2-3.5 mm, tumor: broad-based or endophytic lesions of larger diameter >3.5mm). Animals culled in the first cohort were culled at 7 weeks following tumor development, or if a single tumor interfered with the quality of life before this point. The second cohort animals were monitored and culled when tumor volume occupied >50% of the treated area of the back, or a single tumor interfered with the quality of life. No animals were excluded from the analysis.

### Whole exome sequencing

DNA was extracted from the largest sized tumor for each mouse along with adjacent treated skin, and a kidney using QIAGEN DNeasy Blood and Tissue Kit. Whole exome sequencing was performed by Novogene (Novogene UK). Exome capture was performed with Agilent SureSelect Mouse All Exon Kit and sequenced on the Illumina HiSeq platform. Sequencing reads were trimmed using Trimmomatic(2), aligned to the mm38 reference genome using BWA(3) and duplicate reads were marked using Picard Tools (http://broadinstitute.github.io/picard). Somatic mutations were called using the Varscan 2 pipeline using reads with mapping quality >=20, and tumor and normal coverage >=10(4). Variants within the targeted capture genes were kept for further analysis and were annotated using Variant Effect Predictor(5). Variants present in dbSNP were excluded. Patterns of single nucleotide variation, and insertion and deletions in tumors and skin were analysed using MAFTools in R(6) (version 3.5.1). Oncoplots were generated of the top 20 frequently mutated genes by sex and histology to identify patterns of mutations.

### RNA-seq analysis

Fresh whole DMBA/TPA treated skin from the back and normal skin from the abdomen were collected and DNA and RNA extracted using a QIAGEN AllPrep DNA/RNA extraction kit and a QIAGEN TissueLyser II. RNA sequencing of paired skin samples from 6 mice (3 females, 3 males) was performed by Novogene (Novogene UK). Paired-end sequencing reads were trimmed with Trimmomatic(2) and aligned to the mouse reference genome (GRCm38) using STAR version 2.7.0a(7). Transcript counts were generated by htseq-count(8) using an Ensembl version 99 gtf (Mus_musculus.GRCm38.99.gtf). Differential gene expression analysis was performed using the DESeq2 package(9) (version 1.22.2) in R (version 3.5.1). To identify sex specific autosomal responses analysis was performed on protein coding genes with the X and Y chromosomes removed from analysis. Reads counts of genes were filtered by removing any gene with less than 10 counts across all samples. Analysis of normal and treated skin between sex was performed independently. For pathway enrichment analysis genes that were significantly differentially expressed between sexes in normal or treated skin (FDR p-value < 0.2) were compared against the Hallmark Database using the Molecular Signatures Database v7.0 (Msigdb, Broad Institute, https://www.gsea-msigdb.org/gsea/msigdb/index.jsp). Immune cell populations were inferred from RNA-seq data using ImmuCC(10).

### Histological mouse sample analysis

The histological analysis was performed on the four most prominent lesions per mouse, by diameter, blinded for mouse sex. Tumors were scored as papillomas where the histological features revealed papillary or pedunculated neoplasms of the epithelium with marked acanthosis, papillomatosis and hyperkeratosis surrounding a fibrovascular core, with no evidence of invasion of the dermis. cSCC was defined as a carcinoma of the keratinocytes invading the dermis, and graded: Well differentiated where there was an easily recognizable squamous epithelium, abundant keratinization, apparent intercellular bridges, minimal pleomorphism, and only basally located mitotic figures; Keratoacanthoma when in addition to features of squamous differentiation, the lesion had a marked crateriform appearance limited by squamous lips; Moderately differentiated where only focal keratinization was present and features observed ranged between well and poorly differentiated, with more mitoses across the layers of tumor; and Poorly differentiated where minimal keratinization, marked nuclear atypia, multiple mitoses and scarce squamous differentiation was present. Complete body autopsies were performed, and lymph nodes and organs (except brain) assessed by H&E and pan-keratin stains to confirm metastatic deposits. To measure skin thickness, 10 areas of non-DT exposed and DT exposed skin were examined at 20x, with a Leica Biosystems multiheaded microscope, and superposed layers of epidermis counted. The median number of layers was recorded per site, for n=13 animals. To establish the rate of mitosis, we stained a subset of cSCCs (n =12) and adjacent skin (n =24) with Anti-phospho-Histone H3 (Ser10) (06-570) MilliporeSigma. cSCCs and skin were scored by proportion of positive cells incrementally in 4 categories. To compare the density of infiltrating CD4 and CD8 T cells, we stained cSCCs and adjacent skin with CD4 Antibody (<u>14-9766-82</u>) ThermoFisher / eBioscience and CD8a Antibody (<u>14-0808-82</u>) ThermoFisher / eBioscience.The samples were scored in 5 incremental categories.

### Clinical human sample analysis

Age, sex, immune status and histological grade of human primary cutaneous squamous cell carcinoma were audited from 3 UK NHS hospitals: A: the Christie NHS Foundation Trust, Manchester B: Salford Royal NHS Foundation Trust, Manchester and C: St George’s NHS Foundation Trust, London. Samples from immune-competent and immune-suppressed patients were identified from consecutive, routine pathology reports coded for excised cSCC in a period of 18 months at A and B. An additional search for cSCC from immune-suppressed cSCC was performed by identifying patients in clinical lists of organ transplant recipients/ immunosuppressed patient skin cancer clinics seen at B. Samples from immunocompetent and suppressed patients from C were identified from consecutive multidisciplinary team meetings and from consecutive dermatology-led organ transplant skin cancer clinics. Bowen’s, in situ and rare histological variants of cSCC (SCC with osteoclast◻like giant cells, with sarcomatoid differentiation, pseudovascular/ pseudoangiomatoid SCC, lymphoepithelioma◻like SCC, adenosquamous SCC, spindle◻cell, acantholytic, clear-cell SCCs and epithelioma cuniculatum) were excluded from the audit. Samples were graded by NHS pathologists under routine diagnostic practice according to increasing degrees of histologically aggressive features: keratoacanthoma (KA)/ well differentiated invasive SCC (1), moderately differentiated SCC (2) and poorly differentiated SCC (3). Occasionally, the pathology report stated tumors depicted two distinct areas of differentiation (well-to-moderately differentiated, moderate-to-poorly differentiated), which we graded intermediately (1.5 and 2.5 respectively). The immune status of patients is not routinely available in the pathology report and was retrieved from clinical histories. The collection of clinical variables at B was done under IRAS approval: 216310, REC reference: 16/LO/2098. The collection of data at A and C was approved as an audit on histological grade of cSCC, sex and treatment. Immunosuppressed patients held the following concurrent diagnoses: organ transplant recipients on immunosuppressive medication, white blood cell dyscrasia or systemic cancer treatment with immunotherapy, radiotherapy or chemotherapy in the past 10 years, chronic inflammatory disorders and autoimmune disorders on systemic immunosuppressive medication.

### Data Analysis

Statistical analysis was performed using GraphPad Prism (Version 7.01). Mann-Whitney U test was used to compare significant difference between groups. Human single cell RNA-seq data from a study of squamous cell carcinoma(1) was downloaded from GEO database (GSE144236) and analysed in the Seurat R package(11) following standard pipeline and authors analyses. Cell populations were measured as a percent representation of all cells detected within the sample. Differential expression in the normal tumor-adjacent keratinocytes between sex (female n=3, males n=6) was performed with the FindMarkers function after subsetting samples for normal keratinocytes. Pathway analysis was performed as described above.

## Notes

**Conflicts of interest:** No competing interests

### Competing Interest Statement

The authors have declared no competing interest.

